# An *in vivo* approach to characterize novel variants associated with musculoskeletal disorders

**DOI:** 10.1101/2021.05.27.445925

**Authors:** Jennifer McAdow, Shuo Yang, Tiffany Ou, Gary Huang, Matthew B. Dobbs, Christina A. Gurnett, Aaron N. Johnson

## Abstract

Nemaline Myopathy (NM) is the most common congenital myopathy, characterized by extreme weakness of the respiratory, limb, and facial muscles. Pathogenic variants in *Tropomyosin 2* (*TPM2*), which encodes a skeletal muscle specific actin binding protein essential for sarcomere function, cause a spectrum of musculoskeletal disorders that include NM as well as Cap Myopathy, congenital fiber type disproportion, and distal arthrogrypsosis (DA). *TPM2*-related disorders have not been modeled *in vivo*, so we expressed a series of dominant, pathogenic *TPM2* variants in *Drosophila* embryos and found two variants, K49Del and E122K, significantly affected muscle morphogenesis and muscle function, in part by disrupting myotube guidance. Transient overexpression of K49Del and E122K also disrupted the morphogenesis of immortalized mouse myoblasts *in vitro*, and negatively affected zebrafish muscle development and function *in vivo*. We used our transient overexpression assays in zebrafish to characterize two novel *TPM2* variants that we identified in DA patients (V129A and E139K), and found these variants caused musculoskeletal defects similar to those of the known pathogenic variants. In addition, the severity of musculoskeletal phenotypes in zebrafish expressing *TPM2* variants correlated with the severity of clinical phenotypes observed in DA patients. Our study establishes transient overexpression in zebrafish as an efficient platform to characterize variants of uncertain significance in *TPM2 in vivo*, and suggests that this method can be used to predict the clinical severity of variants associated with DA and congenital myopathies.

## Introduction

Tropomyosins are obligate actin binding proteins that form hetero- and homodimers (Hardeman et al., 2020). Head-to-tail tropomyosin polymers assemble along the length of actin filaments, and in the sarcomere tropomyosin regulates contractility by controlling the ability of thick filament myosin to access actin thin filaments (Squire et al., 2017). To initiate muscle contraction, Ca^2+^ released from the sarcoplasmic reticulum binds to sarcomeric troponin, which alters thin filament confirmation. The intermediate thin filament confirmation allows myosin to contact actin and further displace tropomyosin to drive maximal thin filament sliding and complete contraction (Squire et al., 2017).

Tropomyosin is encoded by four loci in humans (*TPM1, TPM2, TPM3*, and *TPM4*), with *TPM2* and *TPM3* being the predominant skeletal muscle isoforms (Hardeman et al., 2020). Pathogenic *TPM2* variants are causative of congenital skeletal muscles diseases, and much attention has been given toward understanding how *TPM2* variants disrupt sarcomere function. However, tropomyosin also functions outside of the sarcomere to regulate cytoskeletal changes that drive cell migration and cellular metastasis (Bugyi et al., 2010; Lees et al., 2013; Shin et al., 2017). Since skeletal muscle development depends on cytoskeletal dynamics to direct muscle precursor migration (Wang et al., 2018) and myofiber morphogenesis (Williams et al., 2015; Yang et al., 2020), it is distinctly possible that *TPM2* variants adversely affect cytoskeletal dynamics prior to sarcomere assembly which could disrupt overall muscle myogenesis.

Congenital diseases associated with *TPM2* include Nemaline Myopathy (NM) and Cap Myopathy (CM), which are both associated with extreme muscle weakness (hypotonia)(Clarke et al., 2009; Davidson et al., 2013; Mokbel et al., 2013; Ohlsson et al., 2008; Tajsharghi et al., 2007b). The diagnostic features for NM and CM are the presence of nemaline bodies and cap-like structures on muscle biopsy. Pathogenic *TPM2* variants are also causative of congenital fiber type disproportion (CFTD), in which highly oxidative type 1 myofibers are predominant and visibly hypotrophic (Clarke and North, 2003). CFTD patients are also hypotonic.

A fourth congenital disease associated with *TPM2* is Distal Arthrogrypsosis (DA). The heterogeneity of DA clinical phenotypes has necessitated subtype classifications with hierarchical criteria (Bamshad et al., 2009). *TPM2* variants are causative of DA type 1 (DA1)(Sung et al., 2003), which is characterized by contractures of the hands and feet including permanently bent fingers (camptodactyly) and clubfoot (talipes equinovarus) (Bamshad et al., 2009). *TPM2* variants are also associated with DA type 2B (DA2B) (Li et al., 2018; Tajsharghi et al., 2007a), which is characterized by facial abnormalities in addition to contractures of the extremities (Bamshad et al., 2009). DA patients often show hypotonia (Marttila et al., 2014; Mroczek et al., 2017), arguing skeletal muscle dysfunction contributes to the overall disease mechanism.

*TPM2* variants are also causative of Escobar variant of multiple pterygium syndrome (EVMPS)(Marttila et al., 2014; Tajsharghi et al., 2012; Vogt et al., 2020). EVMPS patients show joint contractures similar to those reported for DA patients, but EVMPS is distinguished from DA by the presence of webbing (pterygia) at the neck, elbows, or knees (Morgan et al., 2006). It is important to note that hypotonia often extends to the diaphragm in patients carrying *TPM2* variants, which may require lifelong respiratory intervention (Clarke and North, 2003; Marttila et al., 2014; Ohlsson et al., 2008). The broad spectrum of clinical phenotypes associated with *TPM2* mutations has obscured a clear understanding as to how pathogenic *TPM2* variants disrupt skeletal muscle form and function.

While the *in vivo* disease mechanisms that underlie *TPM2* associated disorders are incompletely understood, the inheritance of *TPM2* congenital diseases follows an autosomal dominant pattern (Bamshad et al., 1994; Mokbel et al., 2013). One notable exception would be the pathogenic variant Q210*, which was shown to be autosomal recessive in a consanguineous family with EVMPS (Monnier et al., 2009; Tajsharghi et al., 2012). Over 30 pathogenic *TPM2* variants have been reported, and the variants themselves show a fairly even distribution along the protein (Fig. 1A)(Marttila et al., 2014). TPM2 is comprised of 7 quasi repeats, each divided into one α-sheet and one β-sheet, with one residue per quasi repeat binding actin (Marston et al., 2013). In addition to the quasi repeats, TPM2 forms a coiled-coil that follows the typical heptad repeat of seven residues, labeled *a-g*, where *b* and *f* residues interact with actin and *g* residues are charged. Surprisingly, only one pathogenic variant changes an actin-binding residue (K128E)(Marttila et al., 2014), while 7 variants cluster to charged *g* positions (Fig. 1B). The molecular genetics of *TPM2*-related disorders argues that pathogenic *TPM2* variants are dominant, gain-of-function mutations that indirectly disrupt tropomyosin-actin interactions.

**Figure 1.**
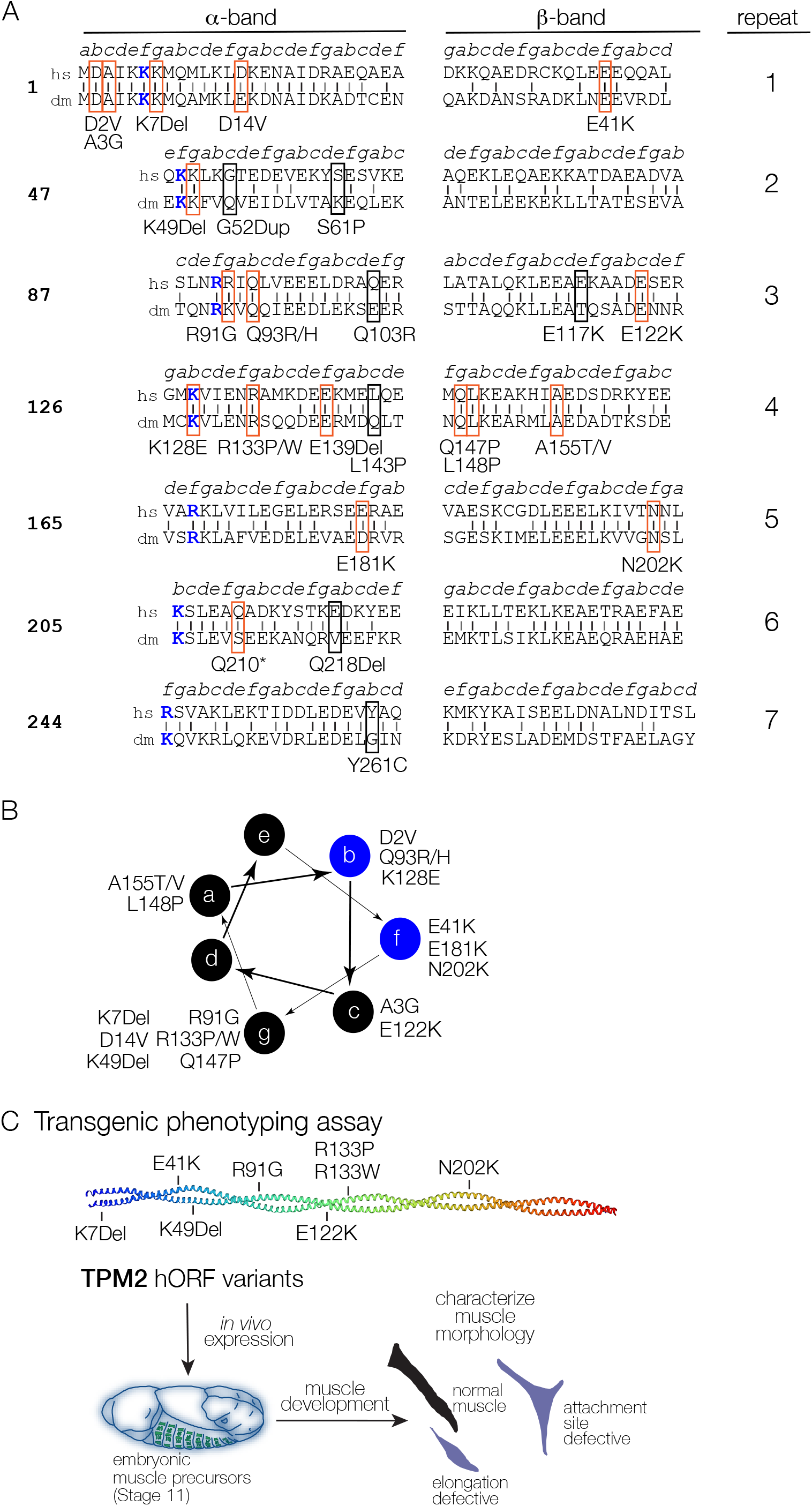
*TPM2* residues associated with pathogenic variants are conserved. (A) TPM2 conservation and sequence structure. TPM2 protein sequence divided into seven quasi repeats and split into α- and β-bands as described in (Marston et al., 2013). Actin-binding residues are colored blue. Vertical black lines show identical residues between the human (hs) and *Drosophila* (dm) proteins; gray lines show similar residues. 25 pathogenic *TPM2* variants in the coding region have been identified (Marttila et al., 2014). Variants affecting conserved residues are boxed in red; non-conserved pathogenic variants are boxed in black. 3 non-coding pathogenic variants have also been reported (not shown). (B) Conserved pathogenic variants partially cluster to a single topographical position. Diagram showing seven amino acids heptad (*a-g*) of the TPM2 coiled coil, as described in (Śliwinska et al., 2018). A subset of *b* and *f* residues binds actin (shown in blue). 7 conserved pathogenic variants mapped to residues in position ‘*g*’. (C) Transgenic expression assay to characterize myogenic defects in muscles expressing pathogenic TPM2 variants. Stable lines encoding TPM2 variants under UAS control were used to express eight conserved pathogenic in specialized embryonic muscle precursors (founder cells); muscle morphology was analyzed at the end of embryogenesis.

Extensive biochemical studies have been used to understand the gain-of-function phenotypes induced by *TPM2* variants. Thin filaments, or even entire muscle fibers, can be reconstituted *in vitro* to assay myosin driven actin motility (Avrova et al., 2018; Borovikov et al., 2020; Borovikov et al., 2017a; Borovikov et al., 2017b; Karpicheva et al., 2020; Marston et al., 2013; Ochala et al., 2010). Reconstituted thin filaments contain actin, tropomyosin, and troponin, such that actin motility can be measured in response to a Ca^2+^ gradient. Actin motility assays have shown that some *TPM2* variants increase Ca^2+^ sensitivity, causing maximum actin motility to be reached at comparatively low Ca^2+^ concentrations, while other variants reduce Ca^2+^ sensitivity (Table 1)(Borovikov et al., 2015; Marston et al., 2013; Marttila et al., 2012; Ochala et al., 2010). The addition of fluorescent probes and proteins to actin motility assays revealed that the Ca^2+^ sensitivity of *TPM2* variants correlates with the ability of troponin and myosin to shift tropomyosin away from actin, and that pathogenic substitutions alter tropomyosin flexibility (Avrova et al., 2018; Borovikov et al., 2020; Borovikov et al., 2017a; Borovikov et al., 2017b; Karpicheva et al., 2020). Since tropomyosin often exists as a heterodimer, *TPM2* variants likely act as gain-of-function mutations by altering Ca^2+^ sensitivity when dimerized with wild-type isoforms (Avrova et al., 2018; Borovikov et al., 2020; Matyushenko et al., 2019). Despite these extensive studies into the biochemical properties of *TPM2* mutations, pathogenic *TPM2* variants have yet to be characterized *in vivo*.

**Table 1.** Properties of pathogenic *TPM2* variants characterized *in vivo*.

| Variant | Diagnosis | Molecular Phenotype | Reference |
| --- | --- | --- | --- |
| K7Del | DA, NM | increased Ca<sup>2+</sup> sensitivity | (Marttila et al., 2014)<br>(Davidson et al., 2013)<br>(Marston et al., 2013)<br>(Mokbel et al., 2013) |
| E41K | DA, NM | reduced Ca<sup>2+</sup> sensitivity | (Avrova et al., 2018)<br>(Marttila et al., 2014)<br>(Marttila et al., 2012)<br>(Ochala et al., 2008)<br>(Tajsharghi et al., 2007b) |
| K49Del | CM | increased Ca<sup>2+</sup> sensitivity<br>reduced actin affinity | (Matyushenko et al., 2019)<br>(Marttila et al., 2012)<br>(Ohlsson et al., 2008) |
| R91G | DA | increased Ca<sup>2+</sup> sensitivity<br>reduced actin affinity | (Robinson et al., 2007)<br>(Sung et al., 2003) |
| E122K | CFTD | unknown | (Marttila et al., 2014)<br>(Tajsharghi et al., 2012) |
| R133P | CFTD | unknown | (Marttila et al., 2014) |
| R133W | DA, EVMPS,<br>NM | reduced Ca<sup>2+</sup> sensitivity | (Vogt et al., 2020)<br>(Borovikov et al., 2020)<br>(Ko et al., 2013)<br>(Ochala et al., 2010)<br>(Tajsharghi et al., 2007a) |
| N202K | CM | unknown | (Jarraya et al., 2012)<br>(Ohlsson et al., 2008) |
| V129A | clubfoot | unknown | this study |
| E139K | DA | unknown | this study |
| A155T | DA | unknown | this study<br>(Jin et al., 2017) |

(CM) Cap Myopathy, (CFTD) Congenital Fiber Type Disproportion, (DA) Distal Arthrogryposis, (EVMPS) Escobar Variant of Multiple Pterygium Syndrome, (NM) Nemaline Myopathy

We set out to model *TPM2* congenital disorders *in vivo*, with the prediction that *TPM2* mutations would adversely affect muscle development and function. *TPM2* has been deleted in mice, and heterozygotes showed compromised lens regeneration (Shibata et al., 2018). However, genome-edited *TPM2* variants have not been reported in any organism. 28 pathogenic *TPM2* variants affecting the coding region have previously been identified, and we used transgenic overexpression in *Drosophila* and zebrafish embryos to study the effects of known variants on embryonic and larval muscle phenotypes. These studies showed that transient overexpression could be useful to model the pathogenic effects of *TPM2* variants, so we expressed three variants that we identified in patients with musculoskeletal disorders in zebrafish. The variants V129A, E139K, and A155T caused phenotypes similar to those we identified for known pathogenic variants, which provided additional evidence that the variants are in fact pathogenic. In addition, the strength of phenotypes from our transient overexpression assays in fish correlated with the severity of patient phenotypes, suggesting our disease model has the power to predict the clinical severity of newly identified variants.

## Results

We set out to model *TPM2* related diseases *in vivo* by characterizing a panel of variants that represent the key characteristics of the spectrum of *TPM2* pathogenic variants. A set of 8 variants in highly conserved residues are causative of the 5 *TPM2* associated disorders, including NM (K7Del, E41K, R133W), CM (K49Del, N202K), CFTD (E122K, R133P), DA (K7Del, E41K, R91G, R133W), and EVMPS (R133W; Table 1). The representative variants are also equally distributed between α-sheets (K7Del, K49Del, R91G, R133P/W) and β-sheets (E41K, E122K, N202K; Fig. 1A). With respect to the TPM2 coiled-coil heptad repeat, pathogenic variants generally cluster to *b*, *f*, and *g* residues (Fig. 1B). The 8 representative variants clustered to the *f* (E41K, N202K) and *g* (K7Del, K49Del, R91G, E122K, R133P) positions, which is consistent with the overall distribution of pathogenic variants along the coiled-coil heptad (Fig. 1B). The variants K7Del, E41K, K49Del, R91G, E122K, R133P, R133W, and N202K are thus a representative collection of *TPM2* mutations causative of congenital disease.

### TPM2 variants disrupt myogenesis

Tropomyosin 2 (Tm2) is the *Drosophila* orthologue of human TPM2, and the two proteins show a high degree of sequence conservation (Fig. 1A). Overexpression studies in *Drosophila* have successfully modeled pathogenic variants in *MYH3* associated with DA (Guo et al., 2020), so we used the binary UAS-GAL4 system to express *Drosophila* Tm2, wild-type human *TPM2*, and the set of 8 human *TPM2* variants in *Drosophila* embryonic muscle precursors (Fig. 1C). For these experiments, UAS constructs were targeted to a common genomic landing site to minimize mRNA expression differences among the variants. The GAL4 lines *slou.Gal4* and *nau.Gal4* direct UAS transgene expression in a subset of muscle precursors (Yang et al., 2020), which allowed us to quantify muscle morphology at single cell resolution (Fig. 1C). Embryonic muscles are named by their position and orientation in the segment, and the Longitudinal Oblique 1 (LO1) muscle shows a striking oblique morphology (Fig. 2A). LO1 muscles that expressed GFP-tagged *TPM2* variants under the control of *slou.Gal4* showed several abnormalities including rounded and generally misshapen morphologies, and attachments to the wrong tendon (Fig. 2A,B). Variant expressing LO1 muscles also failed to develop in the correct position, and were sometimes missing by the end of myogenesis (Fig. 2A,B). The frequency of LO1 muscle phenotypes was higher in muscles that expressed *TPM2* variants than in muscles that expressed wild-type *TPM2* or *Drosophila Tm2, and* LO1 muscles that expressed K49Del showed the highest frequency of muscle defects within the set of 8 pathogenic variants (Fig. 2B). Ventral Oblique 5 (VO5) muscles that expressed GFP-tagged pathogenic variants under the control of *nau.Gal4* were significantly shorter than VO5 muscles that expressed wild-type *TPM2* or *Tm2* (Fig. 2C,D). Among the 8 variants tested, VO5 muscles that expressed E122K showed the strongest phenotype (Fig. 2C,D).

**Figure 2.**
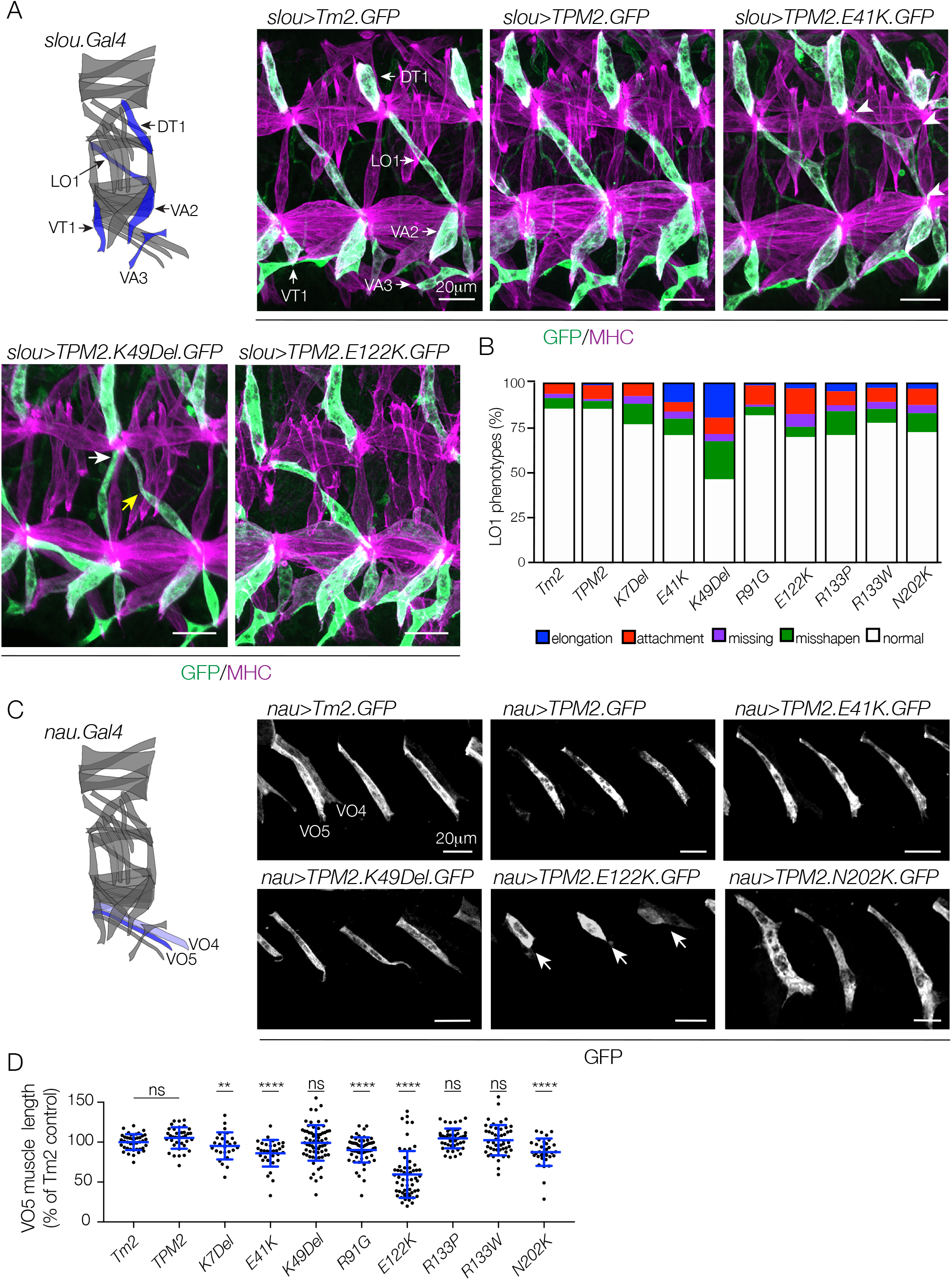
*TPM2* variants disrupt muscle development in *Drosophila*. (A,B) Variants expressed under the control of *slou.Gal4* caused multiple muscle phenotypes. (A) The diagram depicts the thirty body wall muscles in an embryonic hemisegment; *slou.Gal4* expressing muscles are shown in blue [modeled after (Yang et al., 2020)]. Confocal micrographs show Stage 16 embryos that expressed GFP-tagged *Drosophila* Tropomyosin 2 (Tm2), wild-type human TPM2, or pathogenic TPM2 variants (green), co-labeled with Myosin Heavy Chain (MHC, violet). Two hemisegments are shown for each embryo. Variant-expressing LO1 muscles showed multiple phenotypes: rounded muscles (elongation), muscles attached to an incorrect tendon (wrong tendon, white arrows), muscles attached to three tendons (multiple tendons, white arrowheads), muscles absent from a segment (missing), and muscles bent or hook-shaped instead of oblique (misshapen; yellow arrows). (B) Histogram showing phenotypic frequencies (n≥66 muscles/genotype from a minimum of 9 embryos/genotype), quantified by a single blinded protocol. (C,D) Variants expressed under the control of *nau.Gal4* reduced muscle length. (C) The diagram depicts *nau.Gal4* expressing muscles in blue; VO5 muscles show consistent *nau.Gal4* expression and VO4 muscles show occasional expression (light blue). Confocal micrographs show Stage 16 embryos that expressed GFP-tagged transgenes, labeled for GFP. Variant-expressing VO5 muscles were short or rounded, but other parameters of muscle morphology were largely normal. E122K expressing muscles showed the strongest phenotype (white arrows). (D) Dot plot showing VO5 length quantified as % of control; each data point represents a single muscle. Error bars, Standard Error of the Mean (SEM). Significance was determined by student’s t-test versus Tm2-expressing muscles. Scale bars, 20μM.

Our transgenic overexpression studies argue K49Del and E122K act as strong gain-of-function alleles *in vivo*. K49Del deletes a single residue in an α-zone and is associated with CM, while E122K is a substitution in a β-zone and is associated with CFTD (Fig. 1A, Table 1). Surprisingly, neither the position of the variant along the TPM2 protein nor the clinical diagnosis associated with the mutation correlated with the strength of the myogenic phenotype. In addition, the phenotypes we identified in *TPM2* expressing muscles occurred prior to sarcomere assembly, suggesting pathogenic *TPM2* variants affect muscle development and function through sarcomere independent pathways.

### K49Del disrupts myotube guidance

Cellular guidance is a morphogenetic process in which a cell remains spatially fixed, and extends long processes to interact with other cells. During myotube guidance in *Drosophila*, a nascent myotube extends bilateral projections toward tendon cells at the segment border to establish overall muscle morphology. The phenotypes we observed in K49Del expressing LO1 muscles, including short rounded muscles and muscles attached to the wrong tendon, are consistent with defects in myotube guidance (Fig. 2A)(Yang et al., 2020). We have correlated actin dynamics with proper myotube guidance (Williams et al., 2015; Yang et al., 2020), and we suspected that K49Del muscle phenotypes are due to myotube guidance defects.

Live imaging of nascent LO1 myotubes revealed the primary LO1 leading edge initially elongates dorsally, nearly parallel to the dorsal/ventral axis, until it reaches the medial-posterior of the hemisegment (Movie 1). The leading edge then makes a dramatic turn toward the anterior and elongates parallel to the anterior/posterior axis until it reaches a tendon cell at the medial-anterior edge of the hemisegment. The secondary LO1 myotube leading edge elongates a short distance toward ventral-posterior side of the hemisegment where it attaches to a second tendon cell, ultimately giving the LO1 muscle its characteristic oblique morphology.

LO1 myotubes that expressed K49Del showed several unusual behaviors. Some K49Del expressing myotubes would initiate elongation, but both leading edges would retract and show a rounded muscle phenotype at the end of myogenesis (Movie 1). In other examples, the primary and secondary leading edges in K49Del expressing myotubes would elongate appropriately, but the lateral membrane would form a third leading edge and elongate to a third tendon cell (Movie 2). Muscles attached to 3 tendons would presumably be unable to move the exoskeleton correctly. The K49Del variant therefore disrupts myotube guidance, which adversely affects muscle development prior to sarcomere assembly.

### K49Del and E122K inhibit muscle function in vivo

After embryogenesis, *Drosophila* larva develop through three distinct stages (L1-L3), which are characterized by massive muscle hypertrophy (Demontis and Perrimon, 2009). We used *Mef2.Gal4* to express K49Del and E122K in all embryonic and larval muscles, and assayed larval muscle morphology and function. Variant expressing muscles were significantly longer in live L3 larva, which could reflect a reduced contractile state (Fig. 3A,B). Standardized L3 larva locomotion assays have been developed to assess muscle function in insects (Brooks et al., 2016), and we found that K49Del and E122K expressing L3 larva had reduced locomotor activity compared to TPM2 expressing controls (Fig. 3C).

**Figure 3.**
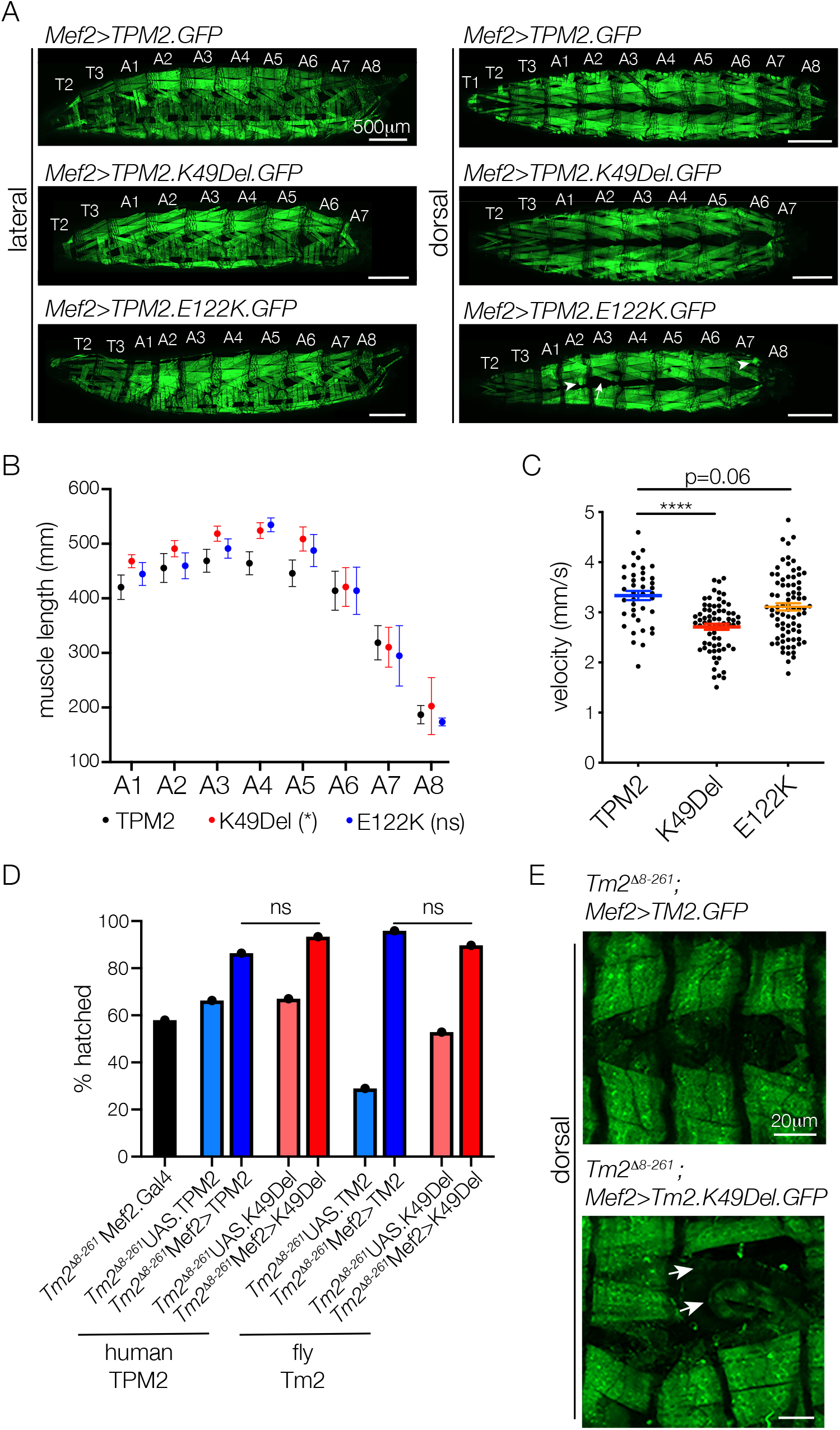
*TPM2* variants disrupt muscle function in *Drosophila*. (A-C) Larval muscles expressing K49Del and E122K have reduced function. (A) Confocal micrographs of live L3 larva that expressed TPM2.GFP, K49Del.GFP, and E122K.GFP under the control of *Mef2.Gal4*. Half of the K49Del.GFP expressing larva completely lacked muscles in segment A8. Muscles were also misshapen (arrowheads) or missing (arrow) in variant expressing L3 larva. Scale bars, 500μM. (B) Dorsal Oblique muscle length. Each symbol represents average muscle length from 8-14 larva/genotype. Variant expressing muscles in segments A3-A6 were longer than controls. (C) Larval locomotion assays. Variant expressing larva moved less than TPM2 expressing controls. (D,E) K49Del disrupted larval muscle morphology in cross-species rescue assays. (D) Rescue assays were performed by crossing *Mef2.Gal4* flies to *UAS.TPM2* or *UAS.Tm2* flies, and measuring hatching rates of the *Mef2>TPM2* and *Mef2>Tm2* embryos. All embryos assayed were homozygous for the *Tm2*^*D8-261*^ allele. Hatching rates were determined for control embryos (*Gal4*-only and *UAS*-only) and rescued embryos (*Mef2>TPM2* and *Mef2>Tm2*). Rescued embryos showed improved hatching rates compared to controls, but hatching rates were similar between wild-type expressing and K49Del expressing embryos. (E) *Tm2*^*D8-261*^ *Mef2>Tm2.K49Del* L1 larva showed muscle defects. Confocal micrographs of live L1 larva. *Tm2*^*D8-261*^ *Mef2>Tm2.K49Del* L1 larva were missing muscles (white arrows), and existing muscles appeared longer than controls. Scale bars, 20μM. Significance was determined by 2-way ANOVA (B) or unpaired student’s t-test (C). (ns) not significant, (*) p<0.05. Error bars, SEM.

We generated UAS constructs encoding untagged TPM2 proteins that were targeted to a common genomic landing site for rescue experiments with a null allele of *Drosophila Tm2*, *Tm2*^*D8-261*^ (Williams et al., 2015). *Tm2*^*D8-261*^ homozygous mutants survive to the end of embryogenesis, but 42.1% failed to hatch presumably due to compromised muscle function (Fig. 3D). We used embryo hatching frequency to assay tropomyosin function, and found wild-type *TPM2* improved *Tm2*^*D8-261*^ embryo hatching (Fig. 3D). However, the hatching frequency was not significantly different between *TPM2* and *K49Del* rescued embryos (Fig. 3D). We repeated the rescue experiments with *Drosophila Tm2*, and found similar results (Fig. 3D), except *Tm2*^*D8-261*^ larva that expressed Tm2.K49Del often lacked muscles, and the remaining Tm2.K49Del expressing muscles appeared longer than controls (Fig. 3E). *Tm2*^*D8-261*^ homozygous animals that hatched showed complete lethality during the L1 stage, but neither wild-type TPM2 nor K49Del significantly rescued *Tm2*^*D8-261*^ L1 lethality. Overall our larval studies confirmed that pathogenic *TPM2* variants inhibit muscle function *in vivo*, but the lack of a complete *Tm2*^*D8-261*^ rescue prompted us to model *TPM2* related diseases in complementary systems.

### TPM2 variants disrupt myotube morphogenesis in vitro

To determine if pathogenic *TPM2* variants also disrupt muscle morphology in vertebrates, we expressed K49Del and E122K in C2C12 cells, which are immortalized mouse myoblasts capable of differentiating into multinucleate myotubes. Under differentiation conditions, C2C12 myoblasts fuse and form nascent myotubes that extensively elongate (Blau et al., 1983). K49Del and E122K proteins mislocalize in C2C12 cells (Abdul-Hussein et al., 2013), but the phenotypes of variant expressing myotubes were not previously characterized. C2C12 cells that expressed untagged, wild-type TPM2 were morphologically similar to control treated cells after 7 days of differentiation, but cells that expressed K49Del and E122K showed significantly reduced myoblast fusion and myotube elongation (Fig. 4A-D). The K49Del and E122K variants therefore disrupt muscle morphogenesis in mammalian cells. Since C2C12 cells develop independent of other musculoskeletal tissues, our studies argue that *TPM2* related disease mechanisms act cell autonomously on developing myofibers.

**Figure 4.**
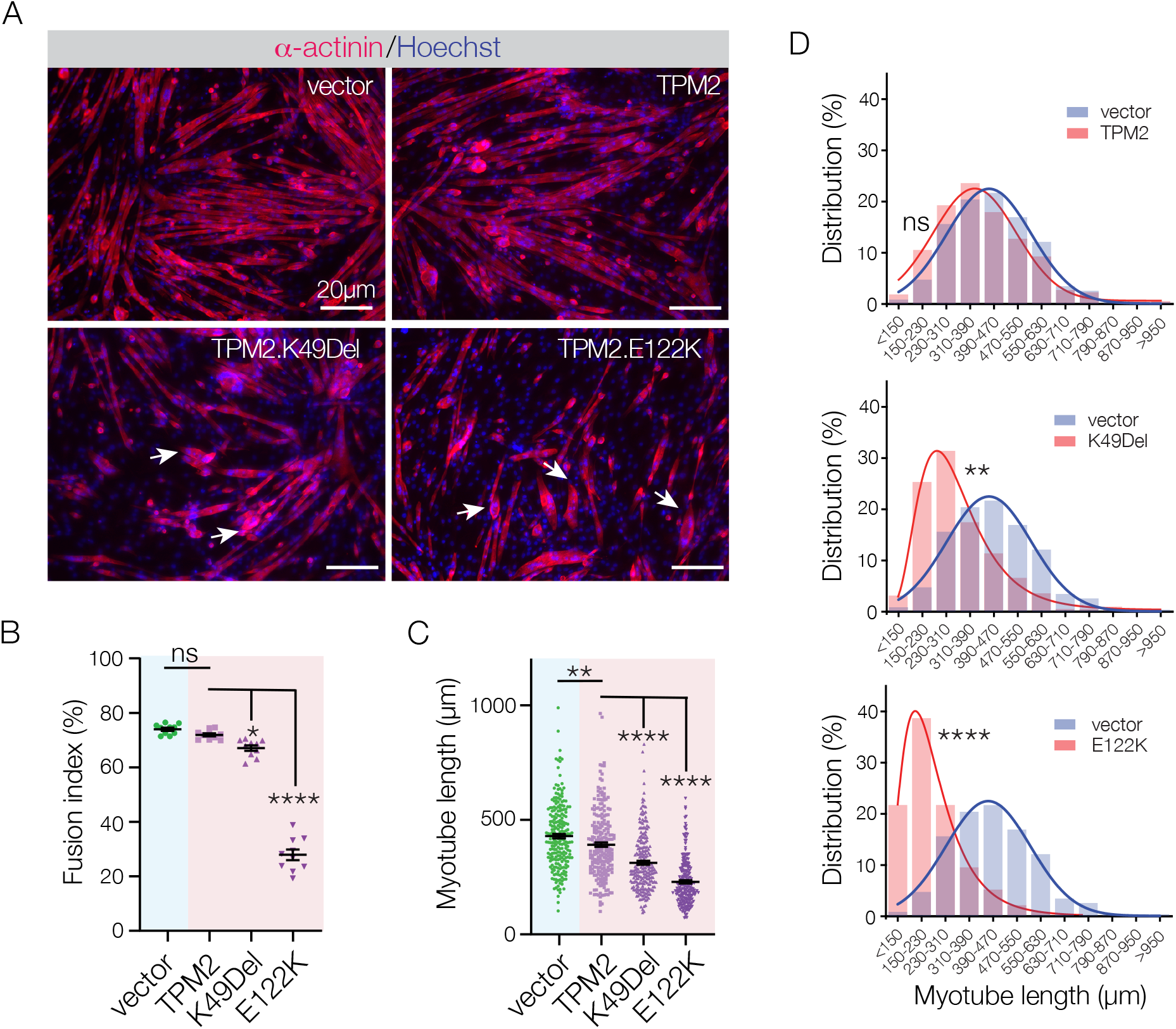
*TPM2* variants disrupt myotube morphogenesis. (A) C2C12 cells transfected with *TPM2* variants showed impaired morphology. Confocal micrographs of cells fixed after 7 days in differentiation media, and labeled for α-actinin (red) to detect differentiated myotubes, and Hoechst to visualize myonuclei. K49Del and E122K expressing myotubes appeared shorter than controls (vector only and wild-type TPM2), and had fewer myonuclei. Variant expressing myotubes were often rounded (arrows). Scale bars, 20μM. (B) Quantification of myoblast fusion. Fusion index represents the number of nuclei in multinucleate myotubes; variant expressing cells fused less than controls. (C) Individual myotubes were measured to quantify cumulative myotube length; variant expressing myotubes were significantly shorter than controls. (D) Myotube length distribution showing the Gaussian distribution fit curve (solid line). Length distribution of variant expressing myotubes skewed toward shorter lengths. Significance was determined by unpaired student’s t-test (B,C) or Mann-Whitney u-test (D). n≥10 imaging fields per treatment. (ns) not significant, **(p<0.01), ****(p<0.0001). Error bars, SEM.

### A transient overexpression assay of variant pathogenicity in zebrafish

Our studies in mammalian cells supported the hypothesis that pathogenic *TPM2* variants disrupt muscle morphogenesis in vertebrates. The K49 and E122 residues are conserved in zebrafish TPM2 (Fig. 5A), and we recently modeled DA2A in zebrafish (Whittle et al., 2020). Genome edited fish heterozygous for the pathogenic variant R672H in *MYH3* showed musculoskeletal abnormalities consistent with joint contractures (Whittle et al., 2020). While the efficiency of genome editing technologies in zebrafish is continuing to improve, the injection of variant-encoding capped mRNAs into fertilized embryos is a well-established tool for rapidly evaluating variant pathogenicity in developing embryos and larva (Jing and Zon, 2011). Though this technique is difficult to use for large transcripts, we took advantage of the comparatively small *TPM2* coding sequence to generate and inject a gradient of mRNA concentrations into one-cell stage embryos and assess muscle morphology at 26hr post fertilization (hpf; Fig. 5B-D).

**Figure 5.**
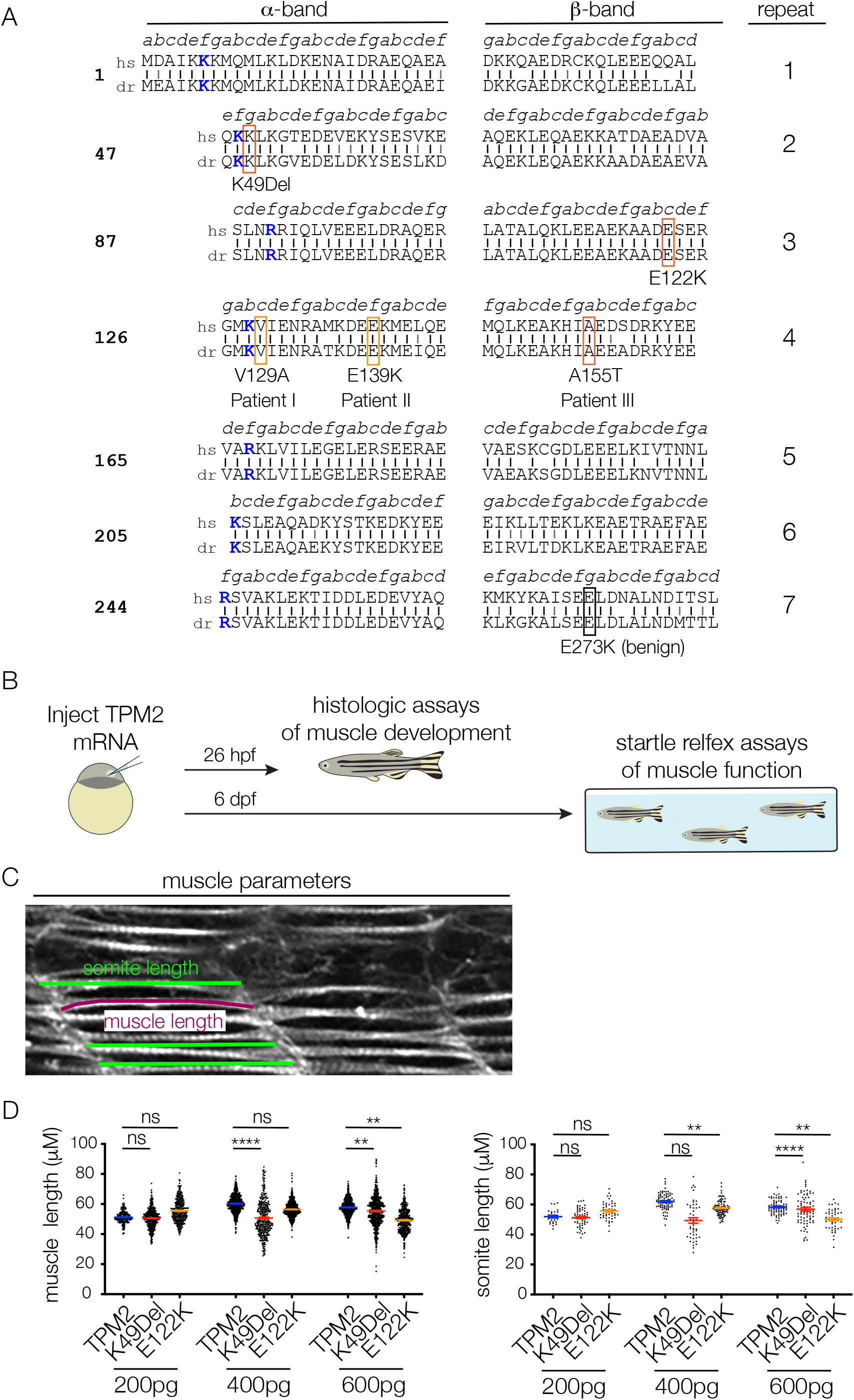
Transient overexpression assays in zebrafish. (A) TPM2 residues associated pathogenic variants are conserved to zebrafish. Vertical black lines show identical residues between the human (hs) and zebrafish (dr) proteins; gray lines show similar residues. Novel variants (V129A and E139K) are boxed in orange; the recurring variant A155T is boxed in red. (B) One cell stage zebrafish embryos were injected with control or variant encoding mRNAs, and raised under standard conditions to 26hr post fertilization (hpf) for histological assays or to 6 days post fertilization (dpf) for startle response locomotor assays. (C) Histologic measurements of 26hpf larva. Individual slow muscle fibers were traced in ImageJ to determine muscle length (magenta line). Somite size was measured 3 times per somite and then averaged to calculate somite length. (D) Pathogenic *TPM2* variants caused dose dependent defects in myofiber length and somite length. A gradient of mRNA levels were injected (200pg, 400pg, and 600pg), and muscle morphology was assessed at each concentration. Significance was determined by unpaired student’s t-test versus wild-type *TPM2* injected fish. Each data point represents an individual muscle fiber or somite. (ns) not significant, (**) p<0.01, (****) p<0.0001. Error bars, SEM.

To validate our transient overexpression assay, we injected embryos with a standardized dose of mRNA (600pg), and quantified muscle and tendon phenotypes in larva that expressed pathogenic or benign variants (Fig. 6A,B). The injection of wild type *TPM2* mRNA or E273K mRNA (rs3180843, LOVD variant 0000446934), which encodes a benign variant identified in a patient with normal muscle function, had negligible effects on myofiber and somite length in the trunk. However, injection of K49Del and E122K encoding mRNAs induced dose dependent defects on musculoskeletal morphology (Fig. 5D, 6A). Larva that expressed K49Del and E122K had significantly shorter slow-twitch myofibers than E273K expressing larva (Fig. 6C). In addition, the slow-twitch myofibers were disorganized, with the myofiber ends often clustered at the somite boundary or even in the center of the somite (Fig. 6A). K49Del expressing larva had significantly fewer slow-twitch myofibers than E273K larva (Fig. 6D), and E122K expressing larva had significantly shorter somite lengths (Fig. 6C). K49Del and E122K did not appear to affect fast-twitch myofiber morphology.

**Figure 6.**
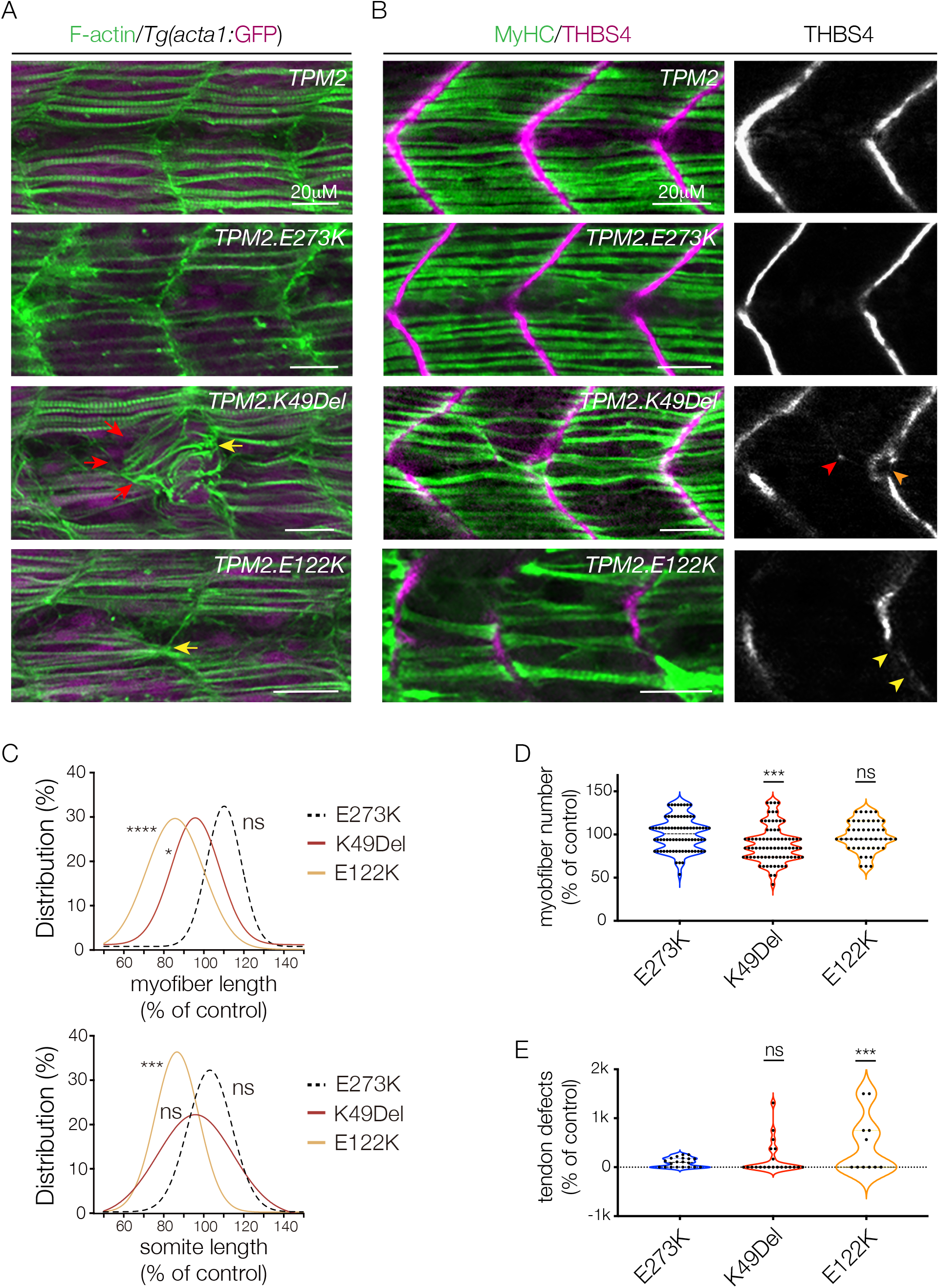
Pathogenic *TPM2* variants disrupt myogenesis in zebrafish. (A) Larva expressing K49Del and E122K showed defects in muscle morphogenesis. Confocal micrographs of 26hpf *Tg(acta1:GFP)* larva injected with *TPM2* RNAs at the one cell stage, labeled for F-actin (green) and GFP (violet). Larva that expressed K49Del and E122K had multiple slow myofiber phenotypes including short fibers (red arrows), and myofibers that clustered to single attachment sites at somite boundaries (yellow arrowheads). Larva that expressed the benign variant E273K had morphologically normal myofibers similar to wild-type controls. (B) Larva that expressed K49Del and E122K showed defects in myosepta morphology. Confocal micrographs of 26hpf larva injected with *TPM2* RNAs at the one cell stage, labeled for slow myofiber Myosin Heavy Chain (MyHC, green) and the myosepta tendon marker Thrombospondin 4 (THBS4, violet). Larva that expressed K49Del and E122K had multiple phenotypes including tendons that developed in the center of the somite (red arrowhead), bifurcated myosepta (orange arrowheads), and myosepta with broken thrombospondin expression (yellow arrowheads). (C) Gaussian distribution fit curves show K49Del and E122K larva had significantly shorter slow myofibers than E273K larva, and E122K had significantly smaller somites. Myofiber length is the average fiber length per somite. n≥48 somites per treatment. (D) Violin plot shows K49Del larva had significantly fewer slow myofibers per somite than E273K larva. n≥48 somites per treatment. (E) Violin plot shows E122K larva had significantly more tendon defects per animal than E273K larva. n≥11 fish per treatment. Scale bars, 20μM. Significance was determined by Mann–Whitney u-test (C), and unpaired student’s t-test (D, E). (ns) not significant, (*) p<0.05, (***) p<0.001, (****) p<0.0001. Error bars, SEM.

Myofibers in the trunk attach to tendons in the myosepta, which are located along the somite boundaries and strongly express the tendon structural protein thrombospondin (THBS4) (Fig. 6B). Larva that expressed K49Del and E122K showed multiple tendon phenotypes that included tendons incorrectly positioned in the center of the somite, bifurcated myosepta, and myosepta with broken or incomplete expression of thrombospondin (Fig. 6B). While both K49Del and E122K caused tendon defects, only E122K larva had a statistically significantly higher frequency of tendon defects than E273K larva (Fig. 6E). Taken together, our studies show that the musculoskeletal phenotypes produced by the transient overexpression of *TPM2* alleles can be used to statistically distinguish pathogenic from benign variants. In addition, our overexpression assays show that K49Del and E122K disrupt musculoskeletal system morphogenesis in zebrafish similar to what we observed in *Drosophila*. These studies provide further evidence that pathogenic *TPM2* variants directly affect myogenesis.

### Novel and recurring TPM2 variants identified in DA patients

Having established a functional assay in zebrafish that can quantitatively identify pathogenic variants in *TPM2*, we decided to evaluate variants of uncertain significance identified in patients with pediatric musculoskeletal disorders treated at Washington University. Patient I, with isolated bilateral clubfoot but no hand contractures, has a heterozygous *TPM2* V129A variant. Patient II and III are unrelated DA1 patients with single heterozygous *TPM2* variants (E139A and A155T; Fig. 7B,C). Patient III also had mild distal lower extremity weakness and fatigue upon running. There was no family history of arthrogryposis or clubfoot, however parents were unavailable for genotyping. Two of the *TPM2* variants, V129A and E139K, had not previously been identified in patients with myopathies or arthrogryposis. The third variant, A155T, was previously identified in a Chinese family with DA1 (Jin et al., 2017). The residues affected by V129A, E139K, and A155T are conserved to zebrafish (Fig. 5A), so we used our transient overexpression assay to provide additional evidence for pathogenicity.

**Figure 7.**
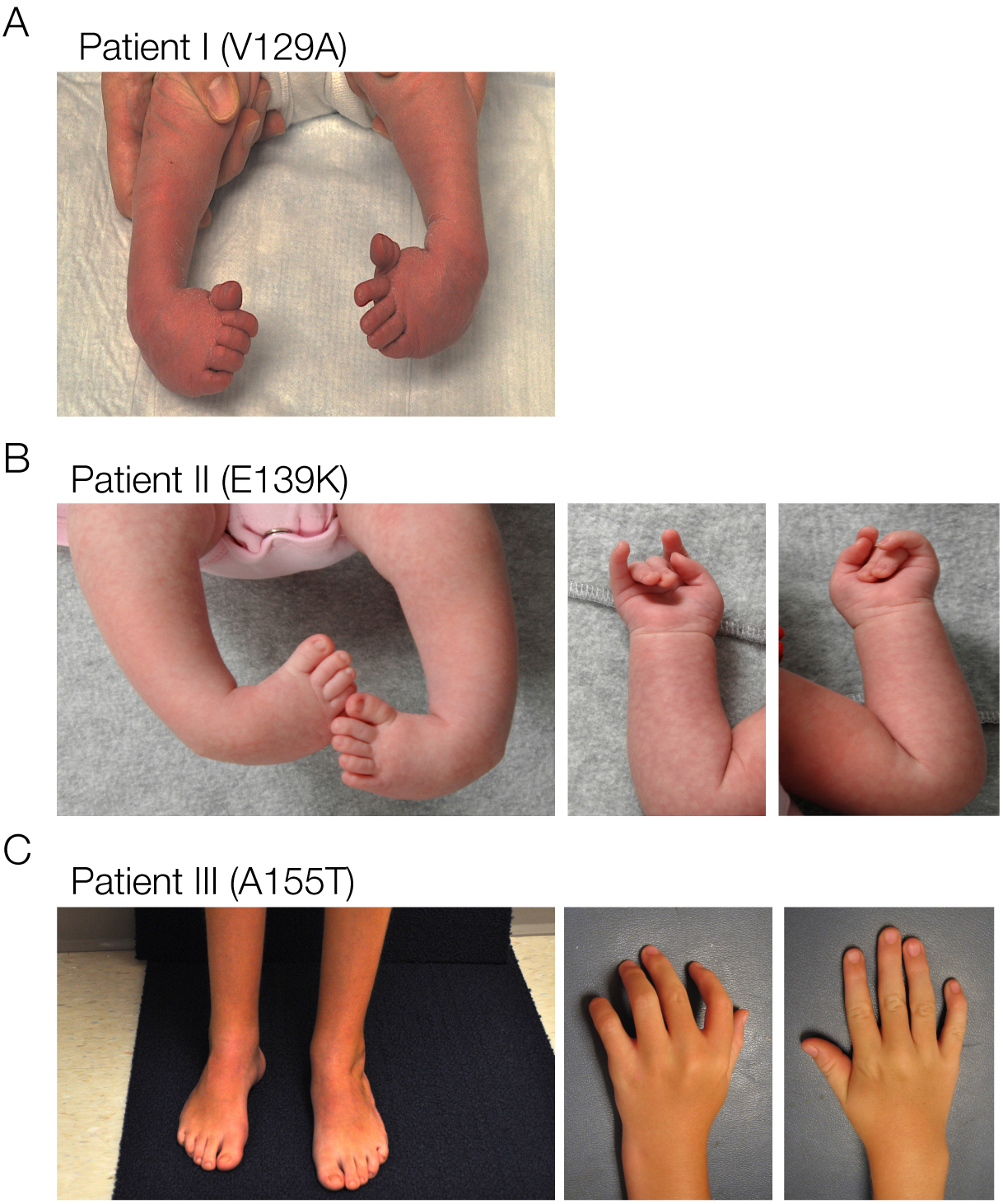
*TPM2* variants were identified in patients with clubfoot and distal arthrogrypsosis. (A) Clinical features of Patient I, diagnosed with bilateral clubfoot (shown here before treatment). (B,C) Clinical features of Patient II and Patient III, diagnosed with DA1. Photos for patient II show clubfoot before treatment, and photos of Patient III show bilateral clubfoot after treatment. Individual III also developed lower extremity weakness as a child. Patient I and Patient II are heterozygous for the novel variants V129A and E139K, respectively; Patient III is heterozygous for the recurring variant A155T.

### Novel TPM2 variants disrupt muscle development and muscle function in zebrafish

Larva that expressed V129A, E139K, and A155T showed defects in musculoskeletal morphogenesis similar to those that expressed K49Del and E122K (Fig. 8A,B). Slow-twitch fibers from A155T expressing larva were significantly shorter and significantly fewer than E273K controls, and A155T somites were significantly smaller (Fig. 8C,D). Larva that expressed V129A and E139K had disorganized myofibers, and V129A myofibers were significantly shorter than controls (Fig. 8A,C). Larva that expressed E139K also showed a significant decrease in somite size (Fig. 8C). In addition, larva that expressed V129A, E139K, and A155T showed a significant increase in the frequency of tendon phenotypes compared to E273K controls (Fig. 8E).

**Figure 8.**
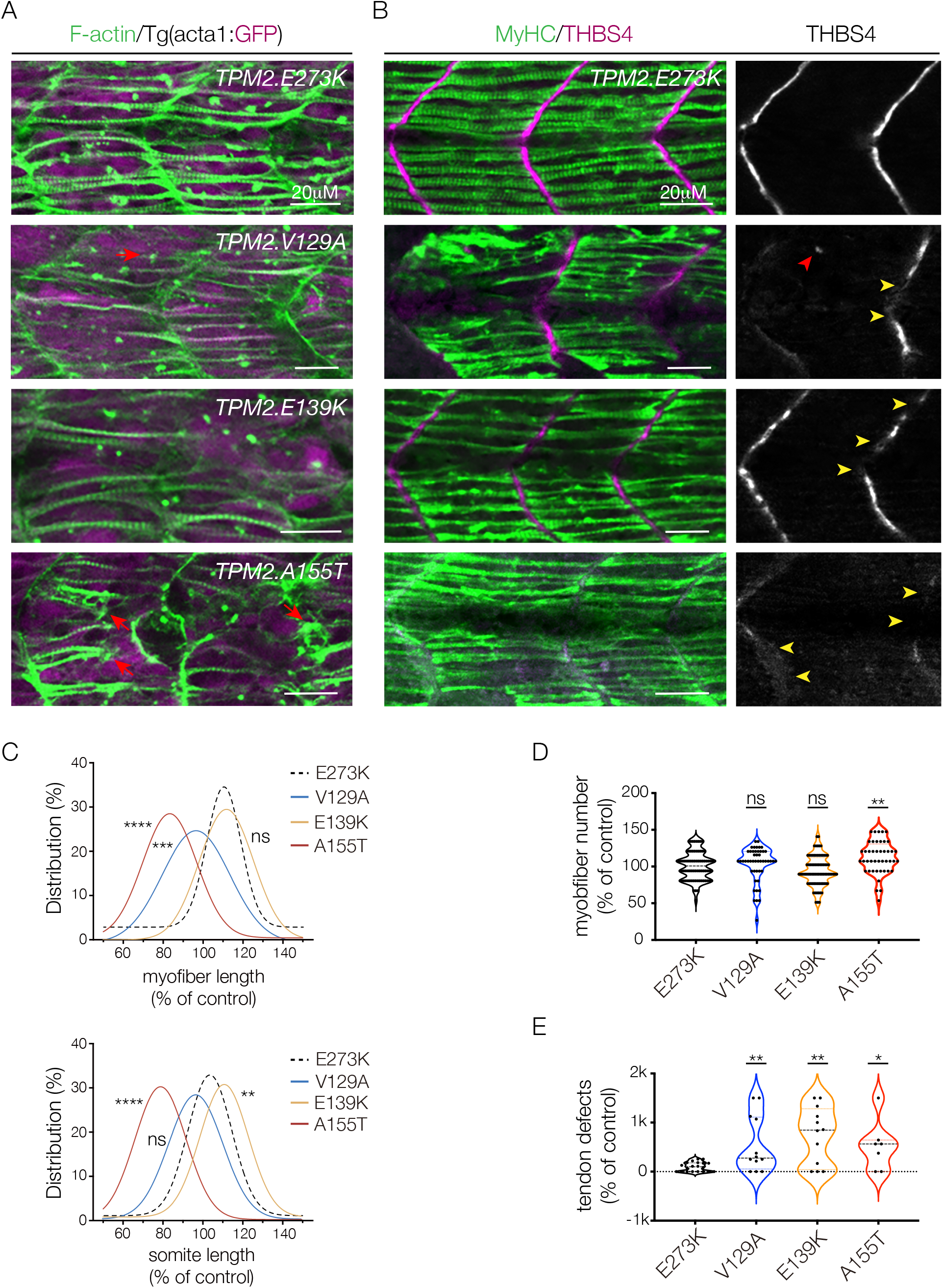
Novel and recurring *TPM2* variants disrupt myogenesis in zebrafish. (A) Larva expressing V129A, E139K, and A155T showed defects in muscle morphogenesis. Confocal micrographs of 26hpf *Tg(acta1:GFP)* larva injected with *TPM2* RNAs at the one cell stage, labeled for F-actin (green) and GFP (violet). Larva that expressed V129A, E139K, and A155T had multiple slow myofiber phenotypes including short fibers (red arrows), and fewer myofibers than larva that expressed the benign variant E273K. (B) Larva that expressed V129A, E139K, and A155T showed defects in myosepta morphology. Confocal micrographs of 26hpf larva injected with *TPM2* RNAs at the one cell stage, labeled for slow myofiber Myosin Heavy Chain (MyHC, green) and the myosepta tendon marker Thrombospondin 4 (THBS4, violet). Larva that expressed V129A, E139K, and A155T had multiple phenotypes including tendons that developed in the center of the somite (red arrowhead), and myosepta with broken thrombospondin expression (yellow arrowheads). (C) Gaussian distribution fit curves show V129 and A155T larva had significantly shorter slow myofibers than E273K larva, and that somite size was significantly different in E139K and A155T larva. Myofiber length is the average fiber length per somite. n≥44 somites per treatment. (D) Violin plot shows A155T larva had significantly fewer slow myofibers per somite than E273K larva. n≥44 somites per treatment. (E) Violin plot shows V129A, E139K, and A155T larva had significantly more tendon defects per animal than E273K larva. n≥12 fish per treatment. Scale bars, 20μM. Significance was determined by Mann–Whitney u-test (C), and unpaired student’s t-test (D, E). (ns) not significant, (*) p<0.05, (***) p<0.001, (****) p<0.0001. Error bars, SEM.

The startle response in zebrafish larva is a well-characterized reflex used to assay motor function, and we previously showed muscle function is compromised in a *MYH3* model of DA using the larval startle response (Whittle et al., 2020). To understand if *TPM2* variants affect muscle function in zebrafish, we injected mRNAs into one-cell stage embryos, and ran automated tracking assays in larva 6 days post-fertilization (dpf) (Fig. 5B, 9A). After a stimulus, the startle response induces a reflexive swim behavior that is quantified by distance swam and the escape velocity. K49Del and E122K expressing larva showed an altered startle response, but surprisingly the swim distance and escape velocity of K49Del and E122K larva were not significantly different than E273K expressing larva (Fig. 9B). However, A155T expressing larva showed significantly reduced swim distance and escape velocity compared to E273K expressing larva, and V129A larva showed significantly reduced escape velocity (Fig. 9B). E139K expressing larva showed an affected startle response, but the differences were not significantly different than E273K expressing larva (Fig. 9B). Nevertheless, the *TPM2* variants we identified in DA patients caused defects in muscle morphogenesis and muscle function, with A155T causing the most severe phenotypes. Taken together, our transient overexpression assays provide additional evidence that the novel *TPM2* variants V129A and E139K are pathogenic, while further confirming A155T is causative of DA.

**Figure 9.**
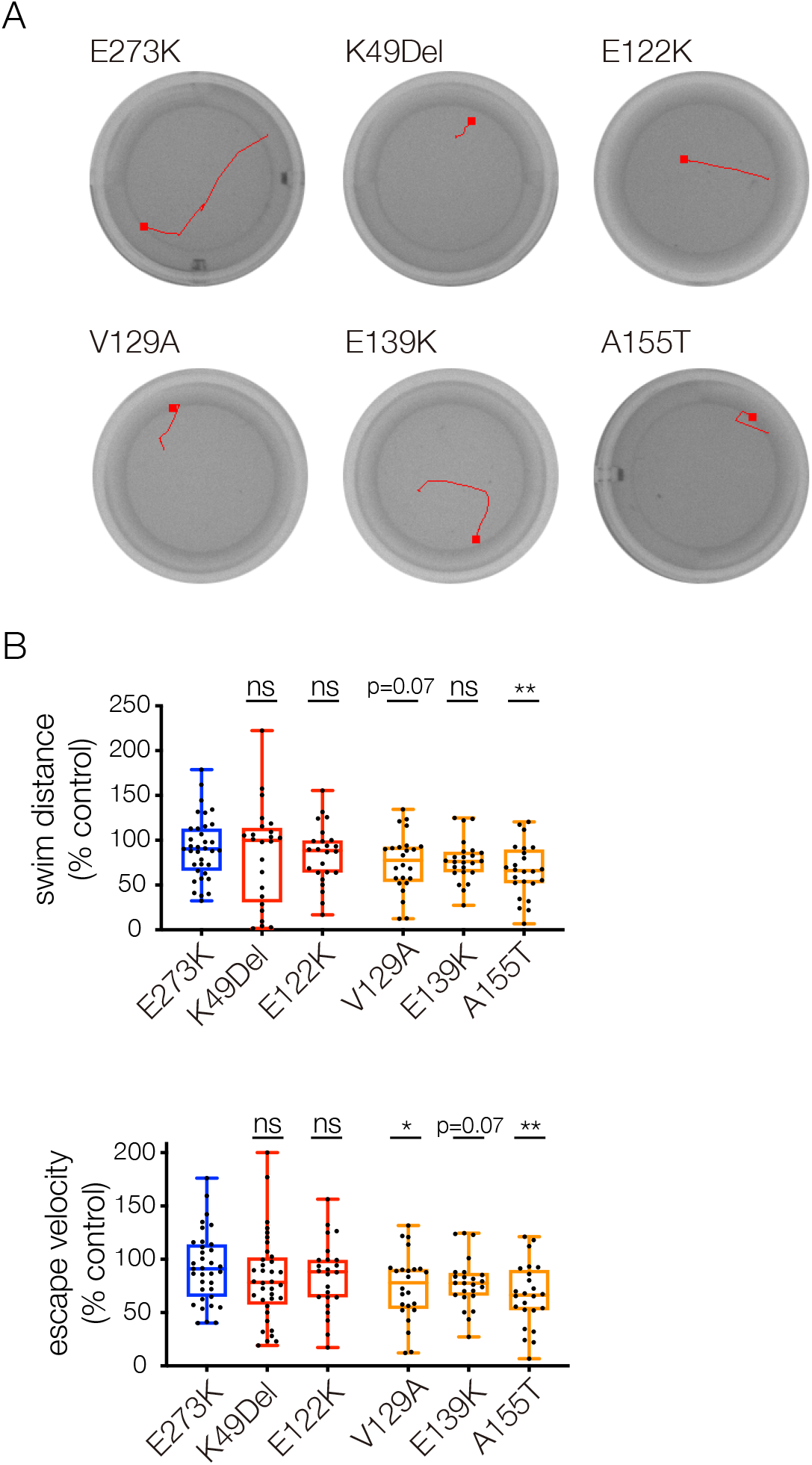
*TPM2* variants affect muscle function in zebrafish. (A) Automated tracking of the startle response in 6dpf larva. Ethovision imaging software tracked larva for 3s after a mechanical stimulus. Red lines show the locomotor path of representative larva. End points are marked by a square. (B) Larva that expressed V129A, E139K, and A155T had reduced startle responses compared to E273K larva. Swim distance and escape velocity of *TPM2* variant injected larva were reported by imaging software, and normalized to wild-type *TPM2* injected larva. Significance was determined by student’s t-test versus E273K larva. Each data point represents an individual fish. (ns) not significant, (*) p<0.05, (**) p<0.01. Error bars, SEM.

## Discussion

To date, over 30 *TPM2* variants have been identified in patients with myopathies and arthrogryposis, but the biochemical properties of only a few variants have been tested. Furthermore, *in vivo* studies investigating the physiological consequences of pathogenic *TPM2* variants were completely lacking. We expressed *TPM2* variants in multiple model systems, and found pathogenic variants disrupt muscle development and muscle function. By focusing on two pathogenic variants, we developed a transient overexpression assay in zebrafish that benchmarked histological phenotypes and functional phenotypes of pathogenic variants against a known benign variant. Clinical sequencing of DA patients identified two novel *TPM2* variants, V129A and E139K, and a recurring variant, A155T, which we tested in our transient overexpression assay. V129A, E139K, and A155T caused musculoskeletal defects similar to those of the known pathogenic variants, and our analyses provide support for pathogenicity of all three variants. Strikingly, A155T was the only variant to cause statistically significant phenotypes in every assay we developed, and the clinical symptoms in the patient with the A155T variant were the most severe among the patients in our study. These results argue that our transient overexpression assay in zebrafish can efficiently characterize variants of uncertain significance identified in patients with musculoskeletal disorders.

Investigations of *TPM2* related disease mechanisms have largely been focused on understanding the role of TPM2 in the sarcomere. Thin filament motility assays uncovered the biochemical properties of TPM2 variants in response to Ca^2+^, and the pathogenic variants tested so far have shown increased as well as reduced Ca^2+^ sensitivity (Table 1). The basis for Ca^2+^ sensitivity is thought to reside in the flexibility or rigidity of the TPM2 dimer, which correlates with the ability of troponin and myosin to shift tropomyosin away from actin (Avrova et al., 2018; Borovikov et al., 2020; Borovikov et al., 2017a; Borovikov et al., 2017b; Karpicheva et al., 2020). Here, we found that *TPM2* variants disrupted muscle morphogenesis prior to sarcomere assembly *in vivo*. Our live imaging in *Drosophila* embryos showed K49Del expressing myotubes had elongation defects and used inappropriate muscle attachment sites to adhere to the exoskeleton (Movies 1,2). We observed similar myotube elongation defects in C2C12 cells that expressed pathogenic *TPM2* variants. Since C2C12 cells develop in the absence of positional cues from other tissues, our studies argue *TPM2* disease mechanisms act cell autonomously to disrupt myofiber morphogenesis prior to sarcomere assembly.

Tropomyosin has well-documented roles outside of the sarcomere in non-muscle cells, most notably during cell migration. Dynamic changes to the cytoskeleton, coupled with changes in the expression of cell adhesion proteins, drive cell migration. Tropomyosins regulate the rate of actin polymerization and depolymerization (Bugyi et al., 2010; Janco et al., 2016; Robaszkiewicz et al., 2016), so it is not surprising that TPM2 and TPM3 are required for single cell as well collective cell migration (Lees et al., 2013; Shin et al., 2017). In vertebrates, myoblasts specified in somites migrate to sites of muscle morphogenesis where they fuse to form myotubes, which in turn elongate and attach to tenocytes (Dennis et al., 1981; Kardon, 1998). During zebrafish myogenesis, myoblasts that give rise to slow- and fast-twitch myofibers are developmentally distinct. Slow myoblasts known as adaxial cells are specified medially, nearest the notochord, and migrate radially to form elongated myotubes on the superficial, outermost region of the somite (Keenan and Currie, 2019). Adaxial cell migration is dependent on cadherin-mediated adhesion, and the slow-twitch region of the myotome in larva that expressed K49Del, E122K, V129A, and A155T (Figs. 6B, 8B) bore a striking resemblance to the slow-twitch myotome of larva that expressed reduced levels of *n-cadherin* (Cortés et al., 2003). It is possible that pathogenic *TPM2* variants disrupt adaxial cell migration in zebrafish, suggesting myoblast migration may be affected in patients with *TPM2* related disorders.

*Drosophila* embryonic myoblasts do not migrate because muscles are specified at the site of myogenesis. However, similar to vertebrates, *Drosophila* myoblasts will fuse to form myotubes that elongate and identify muscle attachment sites (Yang et al., 2020). Myotube elongation and attachment site selection are collectively known as myotube guidance, which is similar to axon guidance in many respects. Cellular guidance is distinct from cell migration because during both myotube guidance and axon guidance, the cell remains spatially fixed but generates long projections to interact with other cells. Myotube guidance depends on regulated cytoskeletal dynamics, particularly of the actin cytoskeleton (Williams et al., 2015; Yang et al., 2020). The *TPM2* muscle morphogenesis defects we have observed in *Drosophila*, zebrafish, and cultured cells are likely the result of improperly regulated actin dynamics in migrating myoblasts or in elongating myotubes that expressed pathogenic variants. DA and amyoplasia (absence of muscle) were often thought to be distinct clinical diagnoses, but amyoplasia was recently reported in a case of congenital DA (Chong et al., 2020). Our studies provide additional support for a model in which DA variants disrupt muscle development. Zebrafish that expressed the DA associated variants E139K and A155T had highly disorganized myofibers that could result in less skeletal muscle mass (Fig. 8A,B). It will be important to develop stable knock in models to characterize the effects of *TPM2* pathogenic variants on myoblast migration, myotube guidance, and long-term muscle homeostasis.

At present, 14 *TPM2* variants have been identified in patients with myopathies and arthrogrypsosis in which the significance of the variant has not been definitively defined (Table 2). The number of *TPM2* variants with uncertain significance is likely to increase because these variants are being identified in patients with isolated clubfoot, which is much more common condition than myopathies or arthrogryposis. The incidence of clubfoot in the United States is 1:1000 live births, but the underlying causes are often unknown. One approach toward understanding isolated clubfoot is to expand clinical sequencing, which will likely uncover novel *TPM2* variants. Phenotypic variability among patients with *TPM2* variants can make genotype-phenotype correlations difficult (Marttila et al., 2014), but the stringency of our benchmarked transient overexpression assay unambiguously distinguishes pathogenic from benign variants. In addition, the assay could be expanded to include other loci associated with myopathies and arthrogrypsosis, which may also uncover new variants that contribute to isolated clubfoot pathogenesis.

**Table 2.** *TPM2* variants of uncertain significance.

| Variant | Clinical significance* | Condition |
| --- | --- | --- |
| D2V | likely pathogenic | not provided |
| Q9P | likely pathogenic | inborn genetic disease |
| S61P | likely pathogenic | NM4 |
| K77R | conflicting interpretations | DA1A |
| R90H** | likely pathogenic | NM |
| Q93H | likely pathogenic | not provided |
| Q103K | likely pathogenic | DA2B |
| E115Del | uncertain | DA1A |
| K128E | uncertain | DA1A |
| H153Q | uncertain | DA1A |
| R178L | uncertain | DA1A |
| D212G | uncertain | DA1A |
| L227V | uncertain | DA1A |
| Q276E | uncertain | DA1A, DA2B, NM |

\*as reported in ClinVar
\*\*all variants showed germline inheritance except R90H (maternal inheritance) (DA1A) Distal Arthrogrypsosis, type 1a (DA2B) Distal Arthrogrypsosis, type 2b (NM) Nemaline Myopathy

One unexpected validation of our transient overexpression assay was that A155T caused the most severe phenotype of any of the variants tested, which also correlated with the clinical severity. The correlation between our laboratory and clinical observations likely reflects the unbiased design of our overexpression assay. Histological phenotypes were quantified by physically measuring individual myofibers and somites in large numbers of larva, and the locomotor phenotypes were quantified automatically by imaging software. We also controlled for day-to-day variation in our assays by normalizing our results to wild-type injected larva each time we injected variants, which allowed us to compare the relative effect of each variant on phenotype severity. One exciting possibility is that our benchmarked transient overexpression assay will have the power to predict the clinical severity of *TPM2* variants, although more variants need to be evaluated to determine the accuracy of our models.

## Acknowledgements

We thank Sharon Amacher for sending *Tg(acta1:GFP)* fish and the Washington University School of Medicine Zebrafish Consortium for providing an outstanding environment to conduct fish work. ANJ was supported by NIH R01AR070299 (NIAMS), and ANJ and CAG were supported by NIH R03HD104065 (NICHD). Research reported in this publication was also funded by the Eunice Kennedy Shriver National Institute of Child Health and Human Development of the National Institutes of Health under Award number 3P50HD103525-01S1 to the Intellectual and Developmental Disabilities Research Center at Washington University and the Washington University Institute of Clinical and Translational Sciences Award number UL1TR002345 from the National Center for Advancing Translational Sciences (NCATS), and R01AR067715-06 (CAG and MBD).

## Author Contributions

Conceptualization: A.N.J., C.A.G, J.M., S.Y.; Methodology: A.N.J., C.A.G, S.Y.; Formal analysis: S.Y., J.M., G.H., C.A.G, A.N.J.; Investigation: S.Y., J.M., G.H., P.H., A.N.J.; Resources: C.A.G., A.N.J.; Data curation: S.Y., J.M., A.N.J.; Writing original draft: A.N.J., C.A.G. ; Visualization: S.Y., A.N.J.; Supervision: S.Y., J.M., A.N.J.; Project administration: A.N.J.; Funding acquisition: C.A.G., A.N.J.

## Materials and Methods

### *Drosophila* genetics

The *TPM2* and *Tm2* transgenic variants were constructed by PCR cloning the *TPM2* ORF (clone HsCD00368588, PlasmID) and the *Tm2* ORF (RE15528, Berkeley *Drosophila* Genome Project) into pEntr (Life Technologies), followed by recombination into destination vector TWG (Drosophila Genome Resource Center) to add a C-terminus GFP tag. TWG clones served as a template to generate variants by site directed mutagenesis as described (Johnson et al., 2013). Tagged and untagged ORFs were PCR subcloned into pUASt.attB using EcoRI/XbaI. All constructs were targeted to the same attP site on chromosome 3L (65B2) using ΦC31 integrase and standard injection methods (Rainbow Transgenic Flies, Inc.). pUASt.attB constructs were fully sequenced prior to injection.

Additional stocks used in this study were *P{GMR40D04-GAL4attP2* (*slou.Gal4*)*, P{GMR57C12-GAL4}attP2* (*nau.Gal4*), *Tm2*^*D8-261*^ (Williams et al., 2015), *P(lacZ-kirre*^*rP298*^) (Nose et al., 1998), and *P(PTT-GC)Tm2*^*ZCL2456*^ (Buszczak et al., 2007). *Cyo, P(wg.lacZ), Cyo, P(twi.Gal4), P(UAS.GFP), TM3, P(ftz.lacZ)*, and *TM3, P(twi.Gal4), P(UAS.GFP)* balancers were used to identify homozygous embryos. Fly stocks were obtained from the Bloomington Stock Center unless otherwise referenced.

### Cell culture

*TPM2* mammalian expression constructs were generated by PCR subcloning variants from pUASt.attB into pCMV-IRES-eGFP (Addgene 78264) using XbaI/EcoR1. C2C12 cells were seeded in 6-well-plate format, grown in standard conditions to 60% confluency in growth medium (10% FBS in DMEM), and transfected with 1μg of DNA per manufacturer’s specifications (Lipofectamine 3000, L3000015, Thermofisher). Empty pCMV-IRES-eGFP was used as a control. Growth media was changed to differentiation media (2% horse serum in DMEM) 24hrs after transfection; cells were differentiated for 7 days and prior to fixation.

### Fish genetics and injections

*Danio rerio* were maintained in accordance with approved institutional protocols under the supervision of the Institutional Animal Care and Use Committee (IACUC) of Washington University, which is fully accredited by the AAALAC. Wild-type zebrafish were from line AB. The *Tg(acta1:GFP)*^*zf13*^ line has been previously described (Higashijima et al., 1997). *TPM2* variants were PCR subcloned from pUASt.attB into pCR2.1 (K202040, Thermofisher), capped RNAs were transcribed with a T7 mMessage mMachine kit (AM1344, Thermofisher), and embryos from natural spawning were injected with up to 600pg RNA in phenol red (P0290, Sigma, 1:6). Injected embryos were maintained at 28.5°C in egg water and collected at 26hr for histology, or at 6dpf for functional assays (feeding protocols began at 4dpf). Control injected larva were collected to normalize each cohort.

### Immunohistochemistry, imaging, and image quantification

#### Drosophila

Dechorionated embryos were fixed in 4% formaldehyde, devitellinated with heptane/methanol, and antibody stained as described (Johnson et al., 2013). Antibodies used were α-Mef2 (1:1000, gift from R. Cripps), α-Myosin Heavy Chain (1:600, Abcam, MAC147), α-GFP (1:600, Torrey Pines Biolabs, TP-401), and α-βgal (1:100, Promega, Z3781). HRP-conjugated secondary antibodies in conjunction with the TSA system (Molecular Probes) were used to detect primary antibodies.

#### C2C12 cells

Differentiated cells were fixed for 15min in 4% PFA, blocked in 5%NGS/PBS, and incubated overnight with α-alpha-actinin (A7811, Sigma, 1:1000). Primary antibodies were visualized with an Alexa-fluor 594 conjugated secondary antibody (115-585-003, Jackson ImmunoResearch Laboratories); myonuclei were visualized with Hoechst (H3570, Thermofisher, 1:1000).

#### Zebrafish

Hand dechorionated larva were fixed in 4% PFA for 1hr and directly stained with Alexa-fluor 555 conjugated phalloidin (A34055, Thermofisher, 1:200) for 2hr room temperature, or blocked in 5%NGS/PBST for 1hr and incubated overnight with THSB4 (Abcam, ab211143, 1:100) and α-Myosin Heavy Chain (F59, Developmental Studies Hybridoma Bank, 1:50). HRP-conjugated secondary antibodies in conjunction with the TSA system (Molecular Probes) were used to detect primary antibodies.

#### Imaging

Embryos and larva were imaged with a Zeiss LSM800 confocal microscope; cells were imaged with an inverted Zeiss AxioObserver. *Drosophila* larva were live-imaged in PBT after 5min exposure to diethyl ether. For time-lapse imaging, dechorionated St12 *Drosophila* embryos were mounted in halocarbon oil and scanned at 2min intervals; dechorionated 3-4 somite zebrafish embryos were mounted in 1.5% low melt agarose supplemented with egg water on a depression slide and scanned at 5min intervals. Control and treated samples were prepared and imaged in parallel where possible, and imaging parameters were maintained between treatment groups. Fluorescent intensity and cell morphology measurements were made with ImageJ software.

### Locomotion and Startle Response Assays

*Drosophila* L3 larva locomotion assays were performed as described (Brooks et al., 2016). For zebrafish larva, the Noldus DanioVision and EthoVision software were used to record and quantify larval movement at 6dpf as described (Whittle et al., 2020). Larva were loaded into the DanioVision in 24-well cell culture plates with egg water at random, and acclimated in the DanioVision box for 5-10 min. The culture plate was automatically tapped after acclimation, and the startle response was recorded for 3s; EthoVision software tracked and recorded fish movement, and reported escape velocity and distance traveled. Statistical analyses were performed only between control and experimental groups assayed on the same day.

### Hatching Assays

0-24hr old embryos were collected on grape agar plates, dechorionated, and genotyped using GFP expression from the *TM3, P(twi.Gal4), P(UAS.GFP)* balancer. Genotyped embryos were transferred to a grape agar plate, incubated for 24hr at 25°C, and then scored for hatching. At least two collections were completed per condition.

### Clinical sequencing

All patients were recruited from St Louis Children’s Hospital or Shriners Hospital St Louis. The institutional review board approved this study and all patients and/or parents provided informed consent. Exome sequencing was performed as described in (Sadler et al., 2020) on a cohort of patients with isolated clubfoot and distal arthrogryposis. Variants were validated by Sanger sequencing.

### Statistics

Statistical analyses were performed with GraphPad Prism 9 software, and significance was determined with the unpaired, one-tailed student’s t-test, one-way ANOVA, or nonparametric tests (for non-Gaussian distributions). Gaussian distribution fit curves were generated with Origin 2019 software. Sample sizes are indicated in the figure legends. Data collection and data analyses were routinely performed by different authors to prevent potential bias. All individuals were included in data analysis.

#### Drosophila

Muscle morphology and size was visualized by Tropomyosin conjugated GFP in hemisegments A2-A8, using 6-10 St16 embryos per genotype. For morphology, muscles were assigned a phenotype (normal, missing, misshapen, elongation defect, attachment site defect), reported as a frequency. Myoblast fusion was quantified by counting the number of lacZ+ myonuclei per hemisegment (A2-A8) in *rP298.nlacZ* embryos. Fusion Index = #lacZ nuclei experimental/#lacZ nuclei control*100.

#### Zebrafish

Methods for measuring musculoskeletal parameters are shown in Figure 5, and largely reflect those reported in (Chagovetz et al., 2019). To control for day-to-day variability in embryo injections, muscle measurements were first normalized to the daily control and then reported as a percent of control.

#### C2C12 cells

Absolute myotube length was used for comparisons among treatment groups. Fusion index = #nuclei in multinucleate myotubes/total nuclei *100. A minimum of 10 fields were quantified per treatment for each parameter.

## Movie Captions

**Movie 1. K49Del expressing myotubes fail to elongate.** LO1 myotubes are pseudocolored blue (WT) or red (K49Del). The K49Del expressing LO1 myotube initiated elongation, but then retracted and failed to reach the muscle attachment site.

**Movie 2. K49Del expressing myotubes develop a third leading edge.** LO1 myotubes are pseudocolored blue (WT) or red (K49Del). The K49Del expressing LO1 myotube initiated elongation with two leading edges, but then developed a third leading edge that appeared to attach to a tendon cell by the end of the movie. Notice a motile cell pinched off of the mutant LO1 and left the filed of view (psuedocolored yellow).

